# Human Astrocytes Exhibit Tumor Microenvironment-, Age-, and Sex-Related Transcriptomic Signatures

**DOI:** 10.1101/2021.02.25.432948

**Authors:** Mitchell C. Krawczyk, Jillian R. Haney, Christine Caneda, Rana R. Khankan, Samuel D. Reyes, Julia W. Chang, Marco Morselli, Harry V. Vinters, Anthony C. Wang, Inma Cobos, Michael J. Gandal, Marvin Bergsneider, Won Kim, Linda M. Liau, William H. Yong, Ali Jalali, Benjamin Deneen, Gerald A. Grant, Gary W. Mathern, Aria Fallah, Ye Zhang

**Affiliations:** Department of Psychiatry and Biobehavioral Sciences, Intellectual and Developmental Disabilities Research Center, Semel Institute for Neuroscience and Human Behavior, David Geffen School of Medicine at the University of California, Los Angeles (UCLA), CA, USA; Brain Research Institute at UCLA; Eli and Edythe Broad Center of Regenerative Medicine and Stem Cell Research at UCLA; Molecular Biology Institute at UCLA; Department of Neurosurgery, David Geffen School of Medicine at UCLA; Department of Molecular, Cell and Developmental Biology, UCLA-DOE Institute for Genomics and Proteomics, Institute for Quantitative and Computational Biosciences - The Collaboratory at UCLA; Department of Pathology and Lab Medicine (Neuropathology) and Department of Neurology, David Geffen School of Medicine at UCLA; Ronald Reagan UCLA Medical Center, Los Angeles, CA, USA; Department of Pathology, Stanford University, Stanford, CA, USA; Program in Neurobehavioral Genetics, Semel Institute, David Geffen School of Medicine; Department of Human Genetics, David Geffen School of Medicine at UCLA; Jonsson Comprehensive Cancer Center at UCLA; Department of Pathology, University of California, Irvine, CA, USA; Department of Neurosurgery, Baylor College of Medicine, Houston, TX, USA; Center for Cell and Gene Therapy, Department of Neuroscience, Department of Neurosurgery, Baylor College of Medicine, Houston, TX, USA; Department of Neurosurgery, Stanford University, Stanford, CA, USA

## Abstract

Astrocytes are dynamic cells with important roles in brain function and neurological disease. There are notable species differences between human astrocytes and commonly used animal models. However, changes of the molecular attributes of human astrocytes across disease states, sex, and age are largely unknown, which is a barrier in understanding human astrocyte biology and its potential involvement in neurological diseases. To better understand the properties of human astrocytes, we acutely purified astrocytes from the cerebral cortices of over 40 humans across various ages, sexes, and disease states. We performed RNA sequencing to generate transcriptomic profiles of these astrocytes and identified genes associated with these biological variables. Here, we identified a robust transcriptomic signature of human astrocytes in the tumor-surrounding microenvironment, including upregulation of proliferation processes, along with downregulation of genes involved in ionic homeostasis and synaptic function, suggesting involvement of peri-tumor astrocytes in tumor-associated neural circuit dysfunction. In aging, we also found downregulation of synaptic regulators and upregulation of markers of astrocyte reactivity, while in maturation we identified changes in ionic transport with implications for calcium signaling. In addition, we identified some of the first transcriptomic evidence of sexual dimorphism in human cortical astrocytes, which has implications for observed sex differences across many neurological disorders. Overall, genes involved in synaptic function exhibited dynamic changes in multiple conditions. This data provides powerful new insights into human astrocyte biology in several biologically relevant states, that will aid in generating novel testable hypotheses about homeostatic and reactive astrocytes in humans.

**Significance Statement:** Astrocytes are an abundant class of cells playing integral roles in the central nervous system. Astrocyte dysfunction is implicated in a variety of human neurological diseases. Yet our knowledge of astrocytes is largely based on mouse studies. Direct knowledge of human astrocyte biology remains limited. Here, we present transcriptomic profiles of human cortical astrocytes, and we identified molecular differences associated with age, sex, and disease state. We found changes suggesting involvement of peritumor astrocytes in tumor-associated neural circuit dysfunction, aging-associated decline in astrocyte-synapse interactions, ionic transport changes with brain maturation, and some of the first evidence of sexual dimorphism in human astrocytes. These data provide necessary insight into human astrocyte biology that will improve our understanding of human disease.

## Introduction

Astrocytes are a major component of the central nervous system. These cells display marked morphological complexity and can drastically change state in response to environmental cues (Faulkner et al., 2004; Chen et al., 2008; Sofroniew and Vinters, 2010; Molofsky et al., 2012; Chung et al., 2015; Nimmerjahn and Bergles, 2015; Ben Haim and Rowitch, 2017; Itoh et al., 2018; Soung and Klein, 2018; Verkhratsky and Nedergaard, 2018; Sofroniew, 2020; Escartin et al., 2021). Though astrocytes were long regarded as passive support cells, studies of murine astrocytes found they have active functions that are critical for the development and function of the central nervous system. For example, astrocyte-secreted factors powerfully induce the formation of functional synapses *in vivo* and *in vitro*, which otherwise largely fails to occur (Banker, 1980; Ullian et al., 2001; Christopherson et al., 2005; Kucukdereli et al., 2011; Allen et al., 2012; Singh et al., 2016; Farhy-Tselnicker et al., 2017; Krencik et al., 2017; Stogsdill et al., 2017; Blanco-Suarez et al., 2018). Astrocytes also shape the activation of glutamate receptors at the synapse through the uptake of glutamate at synapse-associated processes (Rothstein et al., 1996; Huang et al., 2004). In addition to important roles in synapse formation and function, astrocytes contribute to engulfment and elimination of synapses in development (Chung et al., 2013; Tasdemir-Yilmaz and Freeman, 2014; Vainchtein et al., 2018; Lee et al., 2020). There is now a variety of evidence showing that astrocytes help shape circuit functions and behavior (Nedergaard, 1994; Parpura et al., 1994; Halassa et al., 2009; Robel et al., 2015; Papouin et al., 2017; Dowling and Allen, 2018; Mu et al., 2019; Nagai et al., 2019; Huang et al., 2020). Various groups have demonstrated that altering intracellular astrocyte signaling *in vivo* can induce abnormal behavior or correct phenotypic behavior in disease models (Chen et al., 2016; Ma et al., 2016; Ng et al., 2016; Kelley et al., 2018; Yu et al., 2018; Ung et al., 2020; Yu et al., 2020; Nagai et al., 2021). Beyond these roles in synapse development and function, astrocytes provide metabolic support to neurons (Walz and Mukerji, 1988; Pellerin and Magistretti, 1994; Lebon et al., 2002; Itoh et al., 2003), maintain extracellular potassium levels (Kuffler et al., 1966; Olsen and Sontheimer, 2008; Kelley et al., 2018), participate in recycling neurotransmitters (Rothstein et al., 1996), regulate blood flow (Mulligan and MacVicar, 2004; Takano et al., 2006; Attwell et al., 2010), and maintain the integrity of the blood-brain barrier (Abbott et al., 2006). Astrocytes are molecularly and functionally heterogeneous, potentially adapting to diverse roles they play in different brain regions (Tsai et al., 2012; Glasgow et al., 2014; Molofsky et al., 2014; Chai et al., 2017; John Lin et al., 2017; Morel et al., 2017; Miller et al., 2019).

Given their many and varied roles in the CNS, astrocytes are frequently implicated in neurological pathologies (Tian et al., 2005; Yamanaka et al., 2008; Ballas et al., 2009; Lioy et al., 2011; Molofsky et al., 2012; Krencik et al., 2015; Robel et al., 2015; Windrem et al., 2017; Laug et al., 2019). Recently, transcriptomic analysis of neuropsychiatric disease found astrocytic genes included in the gene signatures of autism spectrum disorder, bipolar disorder, and schizophrenia (Gandal et al., 2018a; Gandal et al., 2018b). Astrocyte reactivity is also prominent in several neurodegenerative diseases, including Alzheimer disease and Parkinson disease (Beach et al., 1989). In amyotrophic lateral sclerosis (ALS), reactive astrogliosis occurs around degenerating motor neurons, and this reactivity precedes motor neuron death in the rat SOD1 model of ALS (Howland et al., 2002). Further investigation found that overexpressing GLT1 in astrocytes improved neuronal survival and delayed disease onset in the mSOD1 mouse model of ALS (Guo et al., 2003). However, our understanding of human astrocytes significantly trails our knowledge of murine astrocytes (de Majo et al., 2020). Although the majority of major astrocyte functions appear to be shared between mice and humans, it is still imperative to narrow this gap in knowledge as researchers continue to identify important differences between these species in the CNS. Firstly, human astrocytes are notably larger with more elaborate branching than rodent astrocytes *in vivo* and *in vitro* (Oberheim et al., 2006; Oberheim et al., 2009). At the molecular level, previous characterization of human and mouse astrocyte transcriptomes found many genes specifically expressed by human astrocytes (Zhang et al., 2016; Li et al., 2020). At a functional level, behavioral differences were observed *in vivo* when human glial progenitors were transplanted into mice (Han et al., 2013). Animal studies have produced, and continue to produce, a remarkable body of knowledge concerning the many vital astrocytic functions in the brain (e.g. synapse formation, circuit functions). It is because animal models clearly demonstrate the importance of astrocyte biology that complementary analysis is also required in humans. Given the unique morphological, molecular, and functional properties of human astrocytes, do human astrocytes contribute to the unique capacities of the human brain? Addressing this question requires greater understanding of fundamental human astrocyte biology, including investigation of how astrocytes are impacted across age and development, as well as associations with biological variables such as sex.

Astrocyte biology faces an added layer of complexity considering their significant dynamism in response to insult or injury (Poskanzer and Molofsky, 2018). Astrocytes undergoing reactive astrogliosis in response to a challenge can display stark morphological changes, including hypertrophy and retraction of processes, in addition to a plethora of intracellular alterations (Sofroniew, 2020). Reactivity is observed in many neurological disorders including traumatic brain injury, stroke, epilepsy, and neurodegenerative diseases, and there appears to be disease-specific aspects to this response (Beach et al., 1989; Panickar and Norenberg, 2005; Binder and Steinhauser, 2006; Burda et al., 2016; Liddelow et al., 2017; Yu et al., 2020). This raises the question, what characterizes human astrocyte reactivity, and how does this profile differ in response to varying stimuli? Though animal studies are hugely beneficial in elucidating aspects of human disease, the vast majority of therapeutics developed in mice are not effective in human patients (Kola and Landis, 2004). This highlights the burning need to improve our understanding of human astrocytes in healthy and diseased contexts.

With the advent of improved astrocyte purification methodologies, we can now extract highly pure populations of human astrocytes by targeting the astrocytic cell surface protein HepaCAM using antibodies. By employing an immunopanning technique, we previously published human astrocyte transcriptomes of twelve human cortical samples between the ages of 8 and 63 years old (Zhang et al., 2016). In this study, we acutely purified samples from over 40 patients, which now include astrocytes from healthy and diseased brain regions. For the first time, we are also presenting samples under the age of 8, allowing for analysis of human astrocyte maturation, as well as other biological variables of interest. Here, we describe some of the first transcriptomic data of human astrocytes in the tumor microenvironment, as well as changes in astrocyte gene expression associated with maturation, aging, and sex. Among our findings, we see downregulation of synaptic genes in peritumor astrocytes as well as aging astrocytes.

## Materials and Methods

### HUMAN TISSUE

Human tissue was obtained with informed consent and the approval of the UCLA Institutional Review Board. We obtained tissue primarily from brain surgeries at UCLA to treat epilepsy and tumors, plus one postmortem sample with short postmortem interval (<18 hours). All samples were from the cerebral cortex, primarily from the temporal lobe (n = 31), but several samples came from the frontal (n = 9) or parietal lobes (n = 5), or the insula (n =2). Tissue was immersed in 4° C media (saline or Hibernate-A medium) before transfer to the lab for dissection and dissociation. Six samples were obtained from surgeries offsite, which were shipped overnight in 4°C media for dissection and dissociation in the lab. The final cohort includes 7 peritumor samples, 30 epilepsy samples, and 12 controls totaling 49 samples from 41 patients (see Supplementary Table 1).

### PURIFICATION OF HUMAN ASTROCYTES

Human astrocytes were purified using immunopanning, as described in (Zhang et al., 2016). Briefly, we dissected gray matter from cortical tissue and enzymatically digested the tissue with papain (20 units/mL) for 80 minutes at 34.5°C. We then rinsed the tissue in a protease inhibitor solution. We gently triturated the tissue to generate a single-cell suspension, and we passed the cells over a series of plastic petri dishes that were precoated with antibody. The cell suspension was incubated at room temperature for 10-15 minutes on each dish, which contained anti-CD45 antibody (BD Pharmingen 550539) to deplete microglia, O4 hybridoma to deplete oligodendrocyte precursor cells, GalC hybridoma to deplete oligodendrocytes and myelin, or anti-Thy1 (BD Pharmingen 550402) to deplete neurons. Finally, the astrocyte-enriched cell suspension was incubated for 20 minutes at room temperature on a dish coated with anti-HepaCAM antibody (R&D Systems MAB4108) to bind astrocytes. We washed the bound astrocytes with PBS to remove contaminants, and we immediately harvested RNA by applying 700 μL of TRIzol solution (Thermo Fisher Scientific 15596018). TRIzol solution was then flash frozen in liquid nitrogen and stored at -80°C to await RNA purification. The total time from receiving tissue to storing astrocyte RNA took approximately 4 hours.

### RNA LIBRARY CONSTRUCTION AND SEQUENCING

RNA was extracted from frozen TRIzol using the miRNeasy kit (Qiagen 217004), according to the manufacturer’s protocol. We checked RNA quality with the 2200 TapeStation System (Agilent G2964AA) and the RNA high sensitivity assay (Agilent 5067-5579). All RNA integrity numbers were ≥ 6.5, except RNA samples that were not concentrated enough for accurate measurement. We then used the Nugen Ovation RNAseq System V2 (Nugen 7102-32) to generate cDNA libraries, and we fragmented the cDNA using a Covaris S220 focused-ultrasonicator (Covaris 500217). We amplified and prepared these libraries for sequencing with the NEB Next Ultra RNA Library Prep Kit (New England Biolabs E7530S) and NEBNext multiplex oligos for Illumina (NEB E7335S). We performed end repair and adapter ligation, and we amplified the final libraries using 10 cycles of PCR. The sequencing libraries were verified using the TapeStation D1000 assay (Agilent 5067-5582). Indexed libraries were pooled and sequenced using the Illumina HighSeq 4000 sequencer and obtained 18.9 million ± 1.6 million (mean ± SEM) single end, 50 bp reads.

### READ ALIGNMENT AND QUANTIFICATION

We mapped reads using the STAR package (Dobin et al., 2013) and genome assembly GRCh38 (Ensembl, release 91), and obtained 77.0% ± 5.8% (mean ± standard deviation) uniquely aligned reads in all samples. Reads were counted using the HTSeq package (Anders et al., 2015), and reads were subsequently quantified by reads per kilobase per million (RPKM) using EdgeR-limma packages in R.

### DIFFERENTIAL GENE EXPRESSION ANALYSIS OF DISEASE AND SEX

We analyzed differential gene expression of disease and sex using the DESeq2 package in R (Love et al., 2014). In this analysis, we included all samples and used the following command to create our linear model: ∼ factor(Diagnosis) + Age + factor(Sex) + MicroPC, where Diagnosis was a factor with values [Control, Peritumor, Epilepsy], Age was a numeric value in years, Sex was a factor with values [Male, Female], and MicroPC was numeric value measuring microglia contamination. To calculate the “microPC”, we first determined the gene expression of microglia-specific genes (>10x enriched in microglia vs. astrocytes) in all samples, using the data from (Zhang et al., 2016). Then, we performed PCA analysis using the prcomp function in R on the scaled microglia gene expression, and we took the first principal component (PC1) as a summary measure of microglial gene expression in each sample.

### ANALYSIS OF HUMAN AGING GENES

To identify genes associated with aging astrocytes, we began with genes significantly associated with the Age in the DESeq2 analysis of disease and sex, as described above. In order to separate genes that change in aging (i.e. later life) from genes that change in development (early life), we categorized samples in 3 categories, excluding peritumor samples: 0-21 years old (n = 34), 21-50 years old (n = 3), and 50+ years old (n = 5). We compared younger adults (21-50) to older adults (50+), and we narrowed the gene list to those with an average expression > 0.01 RPKM and 1.5-fold differences in the average expression between groups. This yielded a list of 394 gene entries, 277 of which were protein-coding.

### ANALYSIS OF HUMAN ASTROCYTE MATURATION

We analyzed differential gene expression across astrocyte maturation using a two-step process. First, we performed DESeq2 on samples ≤ 21 years old, excluding peritumor samples (n = 35). Model: ∼ factor(Diagnosis) + Age + factor(Sex) + MicroPC + OligPC + factor(Batch). The “oligPC” was calculated in the same manner as the microPC using gene expression of oligodendrocyte-specific genes identified using data from (Zhang et al., 2016). Finally, we filtered out genes that were 10x enriched in human neurons over human astrocytes, using data from (Zhang et al., 2016). This yielded 1,463 gene entries significantly associated with maturation.

The DESeq2 analysis included samples as young as 7 months old, but we could capture changes from earlier stages in development by using our recently published transcriptomic profiles of fetal human astrocytes (Li et al., 2020). We compared 6 fetal samples with our near-adult human samples between 13-21 years old (excluding peritumor, n = 11). For each sample, gene expression was converted to a percentile, where the most highly expressed gene = 1 and the least expressed gene = 0. Next, we conducted parallel t-tests with a Benjamini-Hochberg correction for multiple tests, performed in GraphPad Prism v8. This generated over 10,000 hits. Finally, we combined the two gene lists using an equal number of genes from each list, (i.e. we filtered results from the second analysis to the 1,463 top hits by p-value to match the first analysis), producing a final list of 2,749 genes associated with human astrocyte maturation.

### MOUSE AGING GENES AND HUMAN COMPARISON

We accessed mouse astrocyte RNAseq data from the BioProject database (www.ncbi.nlm.nih.gov/bioproject), accession no. PRJNA417856 (Clarke et al., 2018). We downloaded FASTQ files of sequenced cortical astrocytes at ages postnatal day 7 (n = 3), 10 weeks (n = 3), and 2 years (n = 2). Reads were aligned with STAR 2.6.0 and genome assembly GRCm38 (Ensembl release 100). All samples had >70% uniquely mapped reads. We used HTSeq to generate counts, and then we determined differential gene expression between the two ages using DESeq2. Model = ∼ factor(Age), where Age is a binary factor.

### GENE ONTOLOGY

To identify patterns in our various gene lists, we used the online platform Metascape (metascape.org, (Zhou et al., 2019)). We input all protein-coding genes from our gene sets, and conducted an enrichment analysis with default settings, with the following adjustments: reference data sets were limited to GO datasets (Molecular Functions, Biological Processes, and Cellular Components), and we supplied a list of background genes. For human analyses, the background gene list consisted of all genes with expression ≥0.05 RPKM in 30+% of our samples. For mouse analyses, the background list consisted of genes with ≥0.05 RPKM in 30+% in the mouse samples from (Clarke et al., 2018).

### RNASCOPE IN SITU HYBRIDIZATION

RNAScope in situ hybridization was performed on fresh frozen human tissue collected from surgeries at UCLA. Tissues were embedded in OCT compound (Fisher Scientific 23-730-571) and sectioned at 30 μm. RNAScope Multiplex Fluorescent V2 Assay (ACDBio 323100) was performed per manufacturer’s protocols. Probes were purchased from ACDBio for the following genes: SLC1A3, C3, VIM, and GFAP. Images were captured using the Zeiss LSM 800 confocal microscope using at equal power and exposure across samples. Photoshop v22.1 was used to equally enhance contrast of the images.

### DATA DEPOSITION

We will deposit all human RNAseq data to the Gene Expression Omnibus before publication.

## Results

### Purification of Human Cortical Astrocytes in Health and Disease

We obtained human cortical tissue from patients undergoing neurological surgery. Our final analysis includes 49 samples from 41 patients, with ages ranging from 7 months to 65 years old. Twelve of these samples were taken from healthy regions of the cortex that were resected due to physical proximity of affected areas. Of these 12 samples, 9 were taken from patients with epilepsy, where it was necessary to resect healthy tissue overlying deep epileptic foci. These resected healthy regions showed no signs of epileptogenicity based on MRI, EEG, and pathological examination. Two additional samples were resected from healthy brain tissue en route to resection of tumors not infiltrating the parenchyma of the brain (e.g., trigeminal nerve schwannomas), and the final sample was taken from a neurologically healthy patient shortly after they died from a heart surgery. From this point forward, we refer to these samples as “controls”. A similar cohort of control human astrocytes were sequenced and analyzed previously, where the authors characterized the baseline characteristics of the human astrocyte transcriptome (Zhang et al., 2016); of note, we used remaining RNA from 7 of these samples in this study. In the current study, we also collected a cohort of samples from brain regions affected by a neurological disorder for comparison with control tissue. Affected samples included in the final analysis fall into two categories of diagnosis: 1) 30 resections from patients with a developmental form of epilepsy, Focal Cortical Dysplasia (FCD), where resected regions participated in epileptic activity; and 2) 7 resections immediately surrounding a brain tumor (referred to here as “peritumor”), including glioblastoma, low grade glioma, and metastatic breast cancer (Supplementary Table 1).

We purified human cortical astrocytes using immunopanning (Fig 1A). We removed white matter and generated a single cell suspension with mechanical and enzymatic digestion. The single cell suspension passes over antibody-coated Petri dishes that bind contaminating cell types with cell type specific antigens. This immunopanning protocol uses anti-CD45 antibodies to pull down myeloid cells (i.e. microglia and macrophages), anti-GalC hybridoma cell supernatant to pull down oligodendrocytes and myelin debris, anti-O4 hybridoma cell supernatant to bind oligodendrocyte precursor cells (OPCs), and anti-Thy1 antibodies to bind neurons. Finally, the enriched suspension passes onto a dish coated in anti-HepaCAM antibodies, a cell-surface glycoprotein enriched in astrocytes. We harvested the astrocyte RNA from this dish and used it to perform RNA sequencing (RNAseq). The sequenced samples show high expression of astrocyte marker genes such as GFAP and SLC1A2, and extremely low expression of neuronal, myeloid, and endothelial genes. There are only slight traces of some oligodendrocyte-lineage marker genes (Fig 1B). Using immunopanning, we obtained RNA highly enriched for human cortical astrocytes in healthy and diseased states for bulk RNAseq (Supplementary Table 2).

**Figure 1.**
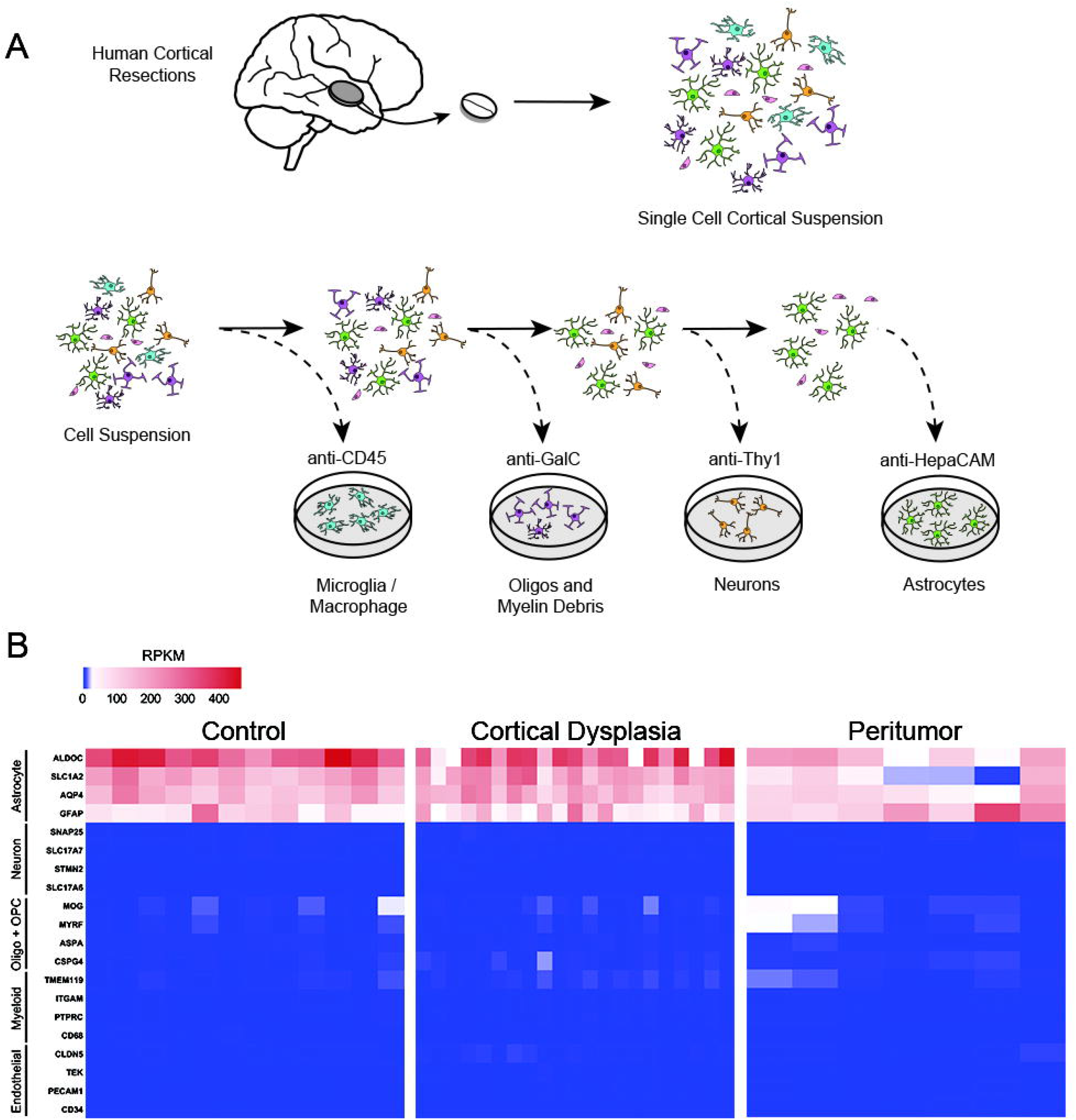
Acute purification of human astrocytes from cerebral cortex. A) Diagram of human astrocyte purification by immunopanning. Surgically resected tissue underwent enzymatic digestion and gentle mechanical digestion to generate a single cell suspension. These cells were passed over a series of plates coated with cell-type-specific antibodies to deplete microglia, oligodendrocyte-lineage cells, and neurons before finally passing to a plate that specifically binds astrocytes using an anti-HepaCAM antibody. B) Heatmaps showing the expression of cell type specific genes in RPKM after RNA sequencing of immunopanned astrocytes. All samples are highly enriched in astrocytic genes (red), with little to no expression of gene markers for neurons, myeloid cells (i.e. microglia or macrophages), oligodendrocyte-lineage cells, or endothelial cells.

### Transcriptomic Signatures of Human Astrocytes in Peritumor Regions and Epilepsy

After sequencing RNA from human astrocytes in epileptic and peritumor regions, we employed differential gene expression analysis to examine their molecular signatures using the analysis package DESeq2 in R. We used a linear model that included variables for diagnosis, sequencing batch, age, and sex. To control for potential variance from low level microglial contamination, we included an additional variable that quantified microglial marker gene expression by performing principal components analysis (PCA) on microglial marker gene expression in our dataset. Including the first principal component in the linear model effectively eliminated significant differential expression of microglial genes.

First, we examined the effect of the brain tumor microenvironment on astrocyte gene expression. Brain tumors drive considerable changes in the surrounding microenvironment, and astrocytes themselves are known to readily change state in response to a variety of stimuli (Raore et al., 2011; Quail and Joyce, 2017). However, transcriptomic changes of peritumor astrocytes in humans have not been reported, to the best of our knowledge. Using our DESeq2 pipeline, we found 3,131 genes differentially expressed in peritumor astrocytes, providing a new resource for elucidating astrocytic changes in the brain tumor microenvironment (Supplementary Table 3). Importantly, many genes related to synaptic function are downregulated in the peritumor region (Fig. 2A-D), such as the glutamate transporters SLC1A2 and SLC1A3, which take up the excitatory neurotransmitter glutamate from the synaptic cleft and maintains excitation-inhibition balance in the brain (Yang et al., 2009). SLC1A2-knockout mice suffer from epileptic seizures and die prematurely (Tanaka et al., 1997). Also downregulated is the gene KCNJ10 encoding the inwardly rectifying potassium channel Kir4.1, which takes up potassium from the extracellular space after neuronal firing, buffers potassium levels in the astrocytic syncytium, and modulates neuronal excitability (Kofuji and Newman, 2004). Patients with mutations in the KCNJ10 gene suffer from seizure disorders (Reichold et al., 2010). A large proportion of human patients with brain tumors also suffer from epileptic seizures, which reduce their quality of life and sometimes cause death (Englot et al., 2016). Our observation of strong reductions of SLC1A2, SLC1A3, and KCNJ10 in peritumor astrocytes suggest potential involvement of astrocytes in tumor-associated seizures and reveal astrocytes as novel potential targets for treating these seizures. A study in a rat model of glioma supports the feasibility of this approach (Sattler et al., 2013). Furthermore, astrocyte secreted molecules that regulate synapse formation and maturation, SPARCL1, CHRDL1, and GPC5 are also downregulated in peritumor astrocytes (Fig. 2B, 2D). Together, these findings suggest decreased support of synapses in the tumor microenvironment that could contribute to dysregulation of synaptic activity and emergent clinical symptoms like seizures. Upregulated genes in peritumor astrocytes include the reactivity-associated genes GFAP, TIMP1 and VIM, suggesting that astrocytes in the peritumor microenvironment are reactive in humans. To assess changes of peritumor astrocytes using an orthogonal approach, we performed *in situ* hybridization with RNAscope. We found that C3, a component of the complement pathway involved in synaptic engulfment and a marker of reactive astrocytes, showed strong astrocytic expression in a sample of peritumor brain, but not in a sample of epileptic brain, as expected based on our RNAseq findings (Fig 2F).

**Figure 2.**
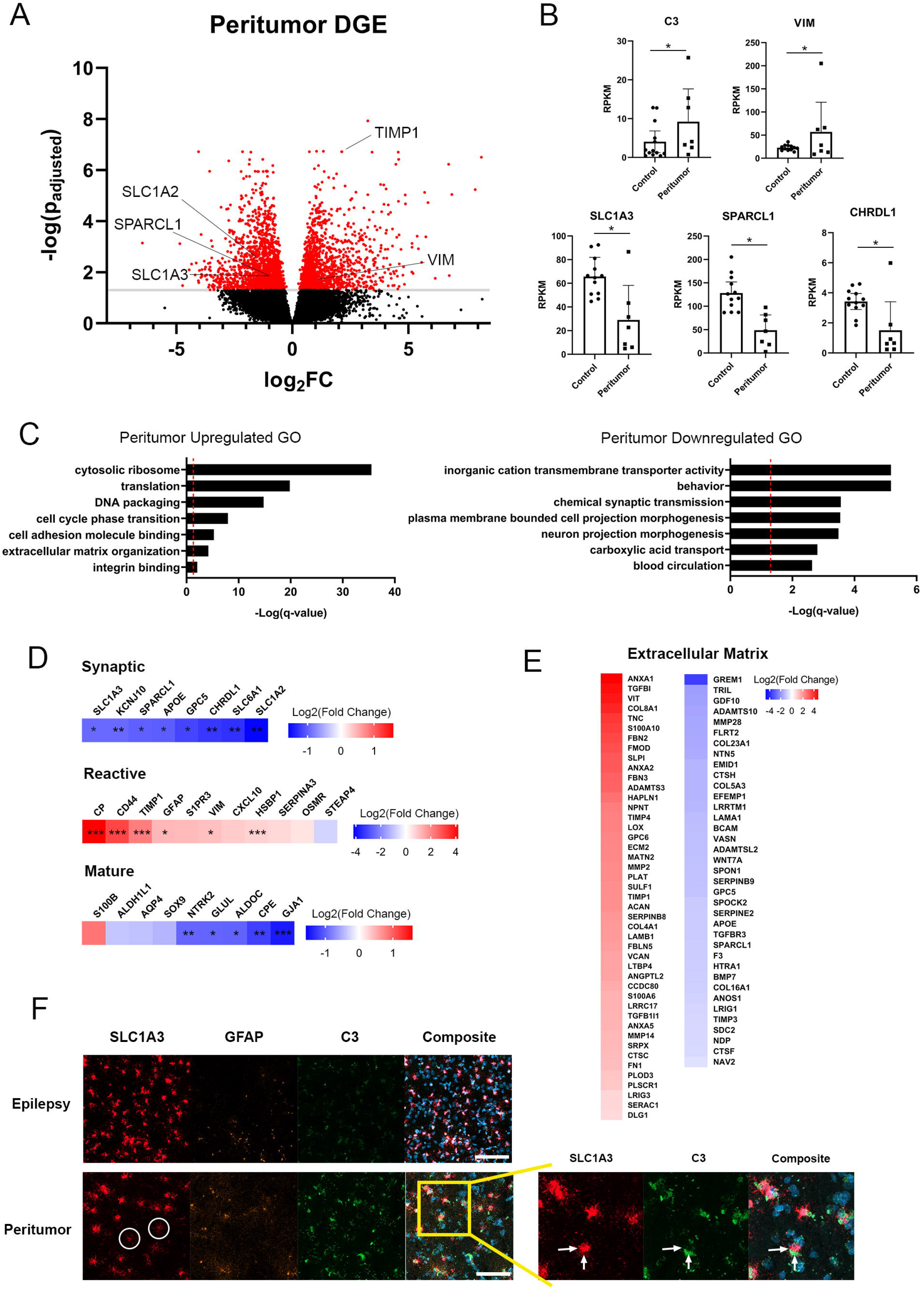
Transcriptomic signature of human astrocytes in the peritumor microenvironment. A) Volcano plot showing differential gene expression in human peritumor astrocytes vs. controls; red = p < 0.05. B) Bar plots of astrocyte genes with changing expression in the peritumor microenvironment. C) Selected gene ontology terms that are significantly enriched in up- (left) and downregulated (right) genes in peritumor astrocytes; dashed lines: p < 0.05. D) Heatmaps of differential gene expression in peritumor astrocytes related to (top) synaptic function, (middle) astrocyte reactivity, and (bottom) mature astrocyte markers. E) Heatmaps of extracellular matrix genes with increased (red) or decreased (blue) expression in peritumor astrocytes, all significant at p < 0.05. F) RNAScope in situ hybridization of human astrocytes, left: Confocal microscopy images of human cortical tissue from patients with epilepsy (top row) or brain tumor (bottom row) stained for SLC1A3, GFAP, and C3 mRNAs. Individual peritumor astrocytes show low SLC1A3 expression (circles). Scale bar = 100 μm. Right: Enhanced magnification demonstrating the distinct localization of SLC1A3 and C3 (arrows).* p < 0.05, ** p < 0.01, *** p < 0.001.

To find larger patterns in the data, we performed pathway analysis with the online tool Metascape to identify gene ontology (GO) terms that are enriched in our gene lists (Supplementary Table 4). Among upregulated genes, we found highly significant enrichment of GO terms related to cell cycle and protein translation, consistent with the presence of proliferative reactive astrocytes in the peritumor region. Meanwhile, downregulated genes were enriched for an array of functional terms related to synaptic function as well as cation transport (Fig 2C), further supporting the hypothesis that peritumor astrocytes are defective in supporting or participating in normal synaptic signaling. Interestingly, both up- and downregulated gene lists are enriched for extracellular matrix genes (Fig 2E), which is notable considering the importance of extracellular remodeling in tumor expansion and migration as well as synaptic plasticity (Nguyen et al., 2020; Winkler et al., 2020). Broadly, we see peritumor astrocytes alter extracellular matrix gene expression while upregulating genes necessary for cell division and translation, and downregulating expression of genes related to synaptic transmission and ionic homeostasis, revealing potential contribution of astrocytes to neural circuit dysfunction associated with brain tumors.

Glioblastoma cells infiltrate surrounding brain tissue. To determine whether our purified astrocytes from the peritumor regions are bona fide astrocytes or infiltrating glioblastoma cells, we examined the expression of a glioblastoma marker gene AVIL (Xie et al., 2020) and did not detect expression in our peritumor astrocyte samples (Supplementary Table 2). Furthermore, we compared gene expression of astrocytes surrounding glioblastoma tissue (infiltrating) and astrocytes surrounding low grade glioma or metastatic tumors (non-infiltrating). Peritumor astrocyte signature genes described above are not more highly expressed by glioblastoma-surrounding astrocytes than non-infiltrating tumor-surrounding astrocytes (Supplementary Table 2). Although we cannot rule out contamination from a small number of infiltrating glioblastoma cells, the gene signatures of peritumor astrocytes are likely from predominantly bona fide astrocytes instead of infiltrating cells.

Next, we examined the transcriptional signature of epilepsy, due to cortical dysplasia. FCD is characterized by abnormalities in neuronal migration during development. Patients with FCD display abnormal radial and/or tangential lamination in a local region of cerebral cortex. More severe cases include dysmorphic neurons, and others also develop large and often multi-nucleated cells called balloon cells (Gaitanis and Donahue, 2013). Our DESeq2 analysis found only 24 protein-coding genes significantly associated with epilepsy, and all but one of those genes (SCN4B) had low expression (an average expression <1 RPKM; see Supplementary Table 5). Samples from epileptiform regions did not separate from controls in PCA, hierarchical clustering, or expression of reactive astrocyte markers (data not shown). Therefore, astrocytes in FCD in humans do not exhibit robust gene expression changes. However, we cannot exclude the possibility that a small subpopulation of astrocytes immediately adjacent to cortical dysplasia lesions have gene expression changes that were undetectable at the population level. Given the lack of robust differences between control samples and epilepsy samples, they were used along with control samples in subsequent analyses.

### Transcriptomic Signatures of Human Astrocyte Maturation

Developing and mature brains have drastically different cognitive capacities, learning potentials, and susceptibilities to disease. Astrocytes are critical for the development of neural circuits, maintenance of homeostasis in adults, and responses and repair in neurological diseases (Huang et al., 2004; Sofroniew and Vinters, 2010; Chung et al., 2013). However, cellular and molecular changes of astrocytes during brain development and maturation in humans are unclear. Previous studies have performed transcriptome profiling of a small number of samples of human astrocytes from fetuses, children ≥8 years old, and adults (Zhang et al., 2016). The gene expression profiles of astrocytes during an important period of development, birth to 8 years, remain unknown. Therefore, molecular knowledge of astrocyte development and maturation in humans is incomplete. Here, we recruited patients throughout development and adulthood (n=16 samples between 0-5 years old; 6 samples between 6-10 years old; 12 samples between 11-17 years old; and 8 adult samples, excluding peritumor; Supplementary Table 1), purified astrocytes, and performed RNA-seq. We analyzed maturation-associated genes based on our new RNAseq data using a linear model (DESeq2 R package, detailed in Methods) and also included fetal data from our previous study after normalizing data from two studies using percentiles (detailed in Methods, (Li et al., 2020)). Our final results find 1509 upregulated genes and 1240 downregulated genes associated with astrocyte maturation across human development (Fig 3C, Supplementary Table 6). To define the time course of astrocyte maturation in humans, we examined the expression trajectory of selected top maturation-associated genes and found that human astrocytes approach mature gene expression profiles around 8 years old (Fig 3A).

**Figure 3.**
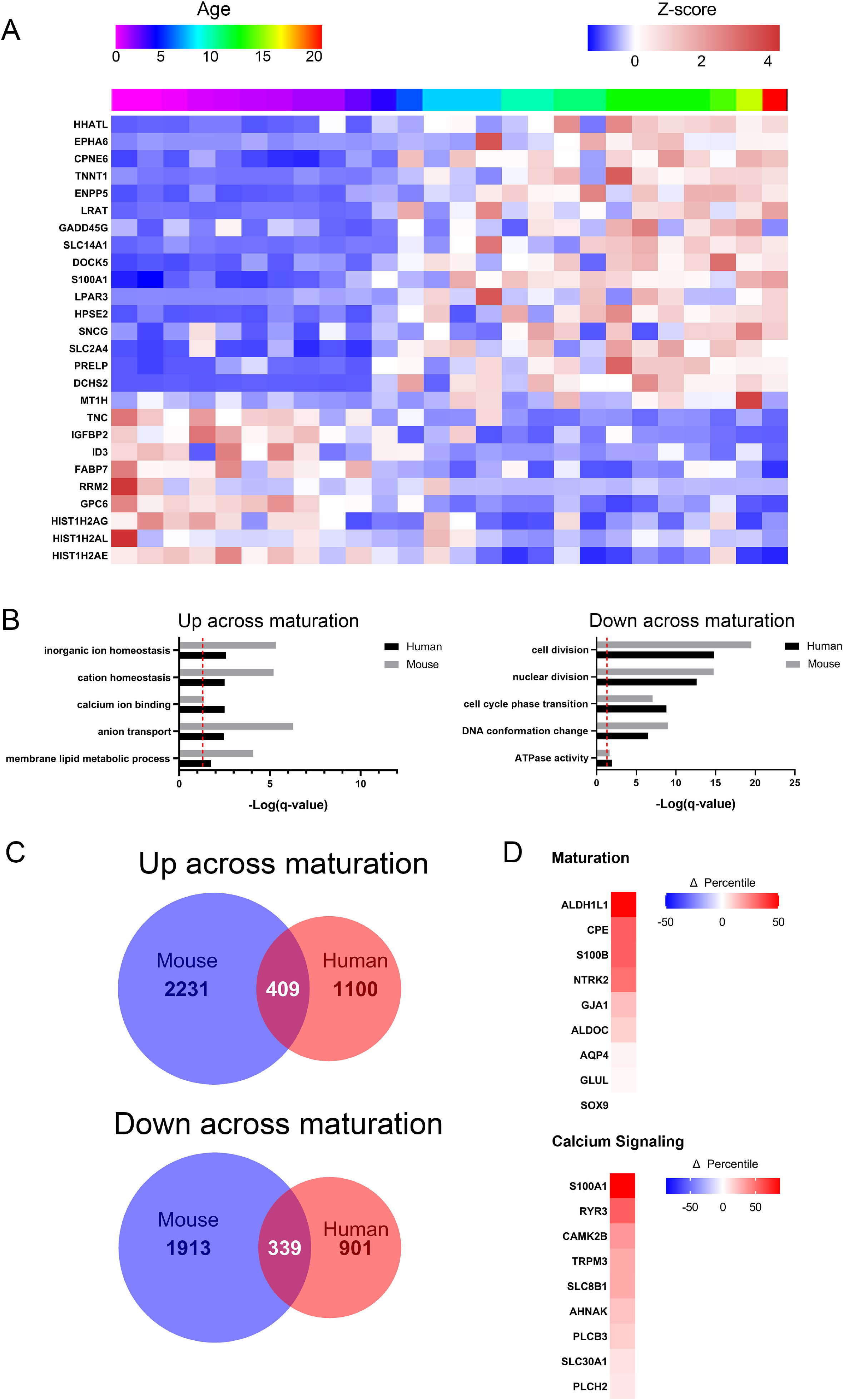
Molecular characterization of human astrocyte maturation. A) Heatmap of representative genes with changing expression across maturation (7 months – 21 years old, n = 26). Astrocytic gene expression approaches the mature pattern around 8 years of age. Plotted as Z-score of gene expression (RPKM); top bar: rainbow index of sample ages. B) Selected gene ontology terms enriched in genes that are up- (left) or downregulated (right) across maturation in both human astrocytes (black bars) and mouse astrocytes (grey bars, from (Clarke et al., 2018)). Dashed lines: p < 0.05. C) Venn diagrams quantifying astrocyte maturation-associated genes that are up- (top) and downregulate (bottom) in humans (red) and mice (blue). D) Top: heatmap of selected astrocyte maturation markers colored by percentile change of RNA expression (e.g. Δ percentile = +100 demonstrates a gene went from the least expressed gene to the most expressed gene) from fetal human astrocytes (Li et al., 2020) to mature human astrocytes (13-21 years old). Bottom: heatmap of selected calcium signaling genes, same quantification as the heatmap above.

To assess the changes associated with astrocyte maturation, we analyzed gene ontology of up- and downregulated astrocyte maturation genes using Metascape (see Supplementary Table 7). Upregulated genes showed significant enrichment for several GO terms related to ion homeostasis, as well as lipid metabolism (Fig 3B). Many of the genes pertaining to ion transport are specifically related to calcium transport and signaling (Fig 3D), which is intriguing given the importance of calcium as a signaling molecule, particularly in astrocytes. Therefore, immature and mature astrocytes may differ in ion transport and calcium signaling, thus altering many downstream signaling pathways that affect both astrocytes and surrounding neurons in development. Downregulated GO terms are almost entirely related to cell cycle and cell division, which is expected to decline throughout development (Fig 3B). We also observe a general upward trend for several astrocyte marker genes that we would also expect to increase across maturation (Fig 3D).

Characterizing the maturation of human astrocytes directly contributes to our understanding of human astrocyte biology and brain development, but most existing knowledge is derived from animal studies. Therefore, it is vital to determine which aspects of human biology are recapitulated by animal models and which are wholly unique. We compared our analysis of human astrocyte maturation with a published study that measured mouse astrocyte gene expression across several ages (Clarke et al., 2018). We compared astrocyte gene expression data from mice at an early developmental age, postnatal day 7 (P7), and a young adult timepoint, 10 weeks, and identified 4,417 genes associated with mouse astrocyte maturation. Most of the mouse and human astrocyte maturation genes we found were not direct orthologues (Fig 3C), and yet both gene lists had remarkably similar patterns based on gene ontology (Fig 3B). Both species downregulate cell division and upregulate ionic transport and calcium signaling genes across maturation. The conservation of these patterns in evolution suggests the importance of these astrocytic developmental changes. Based on this analysis, we find that human and mouse astrocytes share broad outcomes in maturation but differences concerning the exact pattern of molecular changes. While mouse models may not recapitulate every aspect of human biology, our data suggest that the maturation of astrocytes in humans can be accurately modeled in mice.

### Transcriptomic Signatures of Human Astrocyte Aging

Aging is associated with increased risk of cognitive decline and increased susceptibility to neurodegeneration and stroke. Astrocytes are important for maintaining homeostasis of the brain. Yet, aging-associated changes in human astrocytes are largely unknown. Characterizing these changes is the first step in elucidating potential involvement of astrocytes in aging-associated cognitive decline and neurodegeneration and developing astrocyte-targeted treatments. To identify genes with aging-associated expression, we began with genes significantly associated with age in the DESeq2 analysis of all samples that we also used to identify disease-related genes. To identify genes specifically associated with aging (changes after completing maturation) rather than general age (changes across the entire lifespan), we grouped samples into groups by age: 0-20 years old; 21-50 years old; and 50+ years old. From our list of age-associated genes, we extracted genes with average expression >1.5x higher or lower in the 50+ group vs. the 21-50 group. We further filtered by minimum RPKM level of 0.01 to exclude lowly expressed genes. Thus, we identified 394 (277 protein-coding) genes significantly associated with aging (Supplementary Table 8).

As in peritumor astrocytes, we found changes in several synaptic genes when we examined aging-associated gene expression. We find decreased expression of genes mediating astrocyte-synaptic interactions in older astrocytes (Fig. 4A). Most notably, there is a reduction of SLC1A3, a glutamate transporter that clears glutamate from the extracellular space. Under normal conditions, SLC1A3 is a highly expressed core marker of adult astrocytes (Zhang et al., 2016). There is also a decline in CHRDL1, which codes for an astrocyte-secreted factor that drives synapse maturation (Blanco-Suarez et al., 2018). Aging astrocytes also have lower expression of two genes coding for glycoproteins found in the extracellular matrix. The first, CSPG5, shapes neurite growth and localizes around GABAergic and glutamatergic synaptic terminals (Pinter et al., 2020), while the second, OLMF1, binds synaptic proteins such as synaptophysin and AMPA receptors (Nakaya et al., 2013). Declining expression of synaptic genes raises important questions about astrocytic roles in age-related cognitive decline and neurological disease.

**Figure 4.**
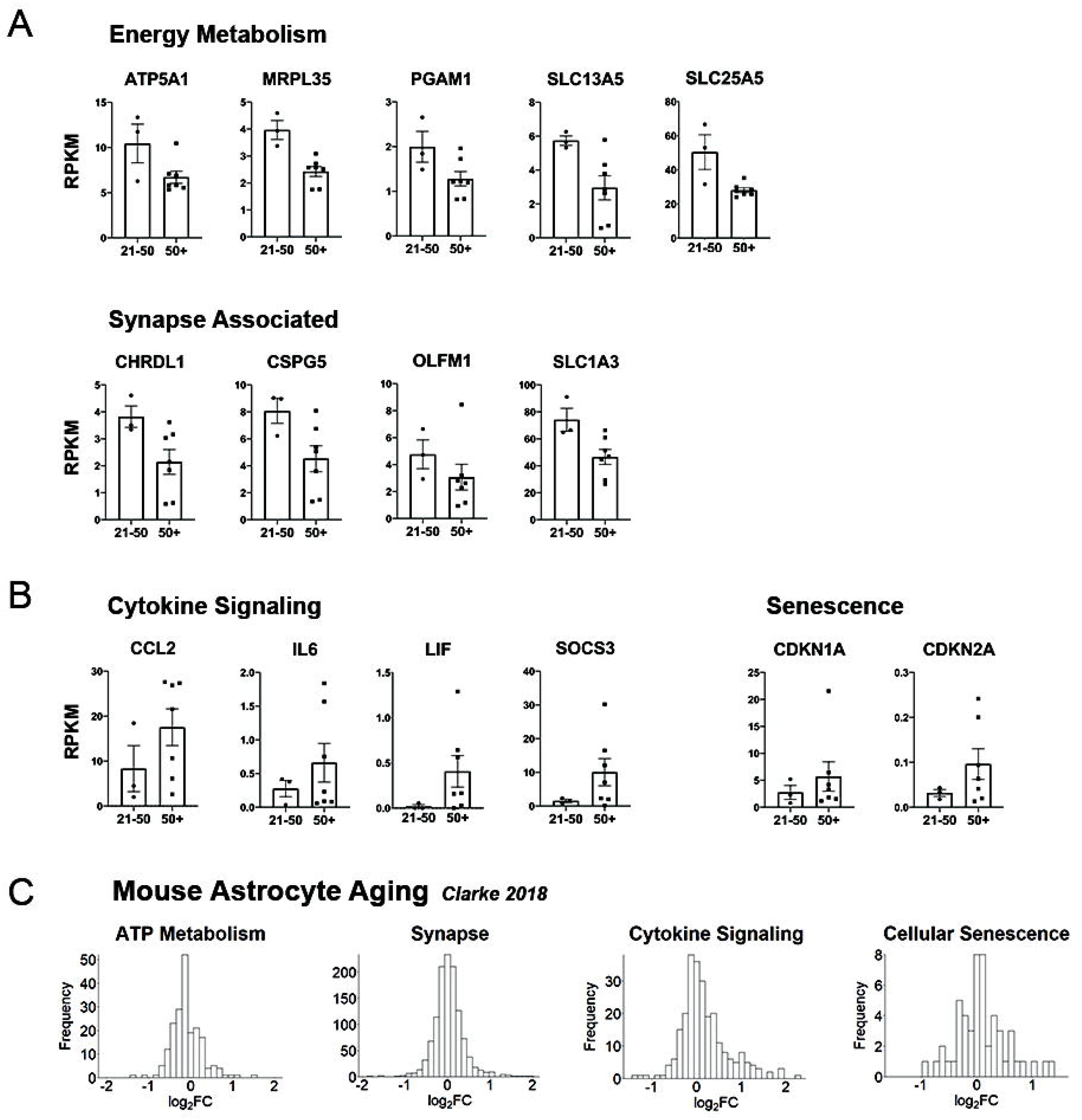
Age-associated genes in human astrocytes. A) Expression of age-associated human astrocyte genes with decreased expression in older adults (50+ years old) compared to younger adults (21-50 years old). B) Age-associated human astrocyte genes with increased expression in older adults compared to younger adults. All genes shown are significantly associated with age, and at least 1.5-fold enriched in younger or older adults. C) Change in RNA expression of mouse astrocytes (10 weeks old vs. 2 years old, from (Clarke et al., 2018)) in various gene ontology categories. Gene lists derived from the following GO annotations, from left to right: GO:0046034, GO:0045202, GO:0019221 and GO:0090398.

Subjects over 50 show additional decreases in genes associated with energy metabolism (Fig. 4A). These include genes involved in mitochondrial generation of ATP such as ATP5A1, an ATP-synthase subunit, MRPL35, a mitochondrial ribosomal component, and SLC25A5, a transporter that carries ATP out of the mitochondria. We also observe lower expression of the glycolytic enzyme PGAM1, and SLC13A5, a citrate transporter. Astrocytes are known to secrete citrate into the extracellular space where citrate has the ability to chelate calcium and magnesium ions, which are important to neuronal NMDA signaling (Westergaard et al., 2017). Together, these gene expression changes are consistent with decreased production of ATP in aging human astrocytes.

Lastly, we also observe an increase of several genes involved in cytokine signaling and senescence. These include the cytokines LIF, IL6, and CCL2, as well as cytokine regulator SOCS3 (Fig. 4B), which are also found in reactive astrocytes in mice (Zamanian et al., 2012). Therefore, these changes suggest that aging astrocytes may exhibit altered energy metabolism, differences in interactions with neuronal synapses and increased cytokine signaling.

To assess the similarity between human and mouse astrocyte aging, we returned to the mouse aging dataset from (Clarke et al., 2018). They reported a list of 58 age-associated genes in mouse cortical astrocytes. We wanted to determine whether mice recapitulated human changes related to metabolism, synapses, cytokines and senescence. Due to the short length of the mouse gene list, we sought to identify trends in the whole dataset. To do so, we first chose relevant GO terms that captured the trends we observed in humans (“ATP metabolic process”, “synapse”, “cytokine-mediated signaling pathway”, and “cellular senescence”). Using these gene lists, we plotted differences in gene expression of all genes between the ten-week-old and two-year-old mice (Fig. 4C). There is a prominent right skew in cytokine signaling genes, and we observe a slight left shift in ATP metabolism genes, suggesting mouse astrocytes also show signs of upregulating cytokine signaling genes while downregulating metabolic genes in old age. The distributions for synaptic and senescence genes were highly symmetrical, suggesting no broad trends associated with age. However, mouse astrocytes could show important changes in smaller subsets of synaptic or senescence genes upon further analysis. In total, we see evidence that mouse astrocytes share metabolic and cytokine features of human astrocyte aging.

### Sexually Dimorphic Gene Signature of Human Astrocytes

Female and male brains differ in their susceptibility to neurological disorders. For example, autism spectrum disorder and Parkinson disease are more prevalent in men than in women (Werling and Geschwind, 2013; Gillies et al., 2014), whereas multiple sclerosis, Alzheimer disease, and anxiety disorder are more prevalent in women than in men (Seshadri et al., 1997; McLean et al., 2011; Westerlind et al., 2014). The cellular and molecular mechanisms underlying sexually dimorphic susceptibility to neurological and psychiatric disorders are largely unknown. While understanding of sexually dimorphic properties of other brain cells, particularly microglia, is increasing, sexual dimorphism in human astrocytes has not yet been reported. In our differential gene expression analysis of all 49 samples, we found 105 genes (40 protein coding) with expression levels significantly associated with sex (Fig. 5, Supplementary Table 9). This gene list represents the first evidence of sexual dimorphism in human cortical astrocytes, to the best of our knowledge. Several of these genes are transcription factors (POU5F1B, HOXC10), some of which are located on sex chromosomes (ZFY, ZFX). Females had higher expression of genes with epigenetic functions, such as demethylases KDM5C and KDM6A, and methyl transferase TRMT6. We also observed a diverse array of differentially expressed genes whose protein products are located in the plasma membrane, though their functions in astrocytes remain mysterious (TMEM176B, TMEM143, CD99). Together, this data represents the first evidence that human astrocytes display a subtle sexual dimorphism at the molecular level.

**Figure 5.**
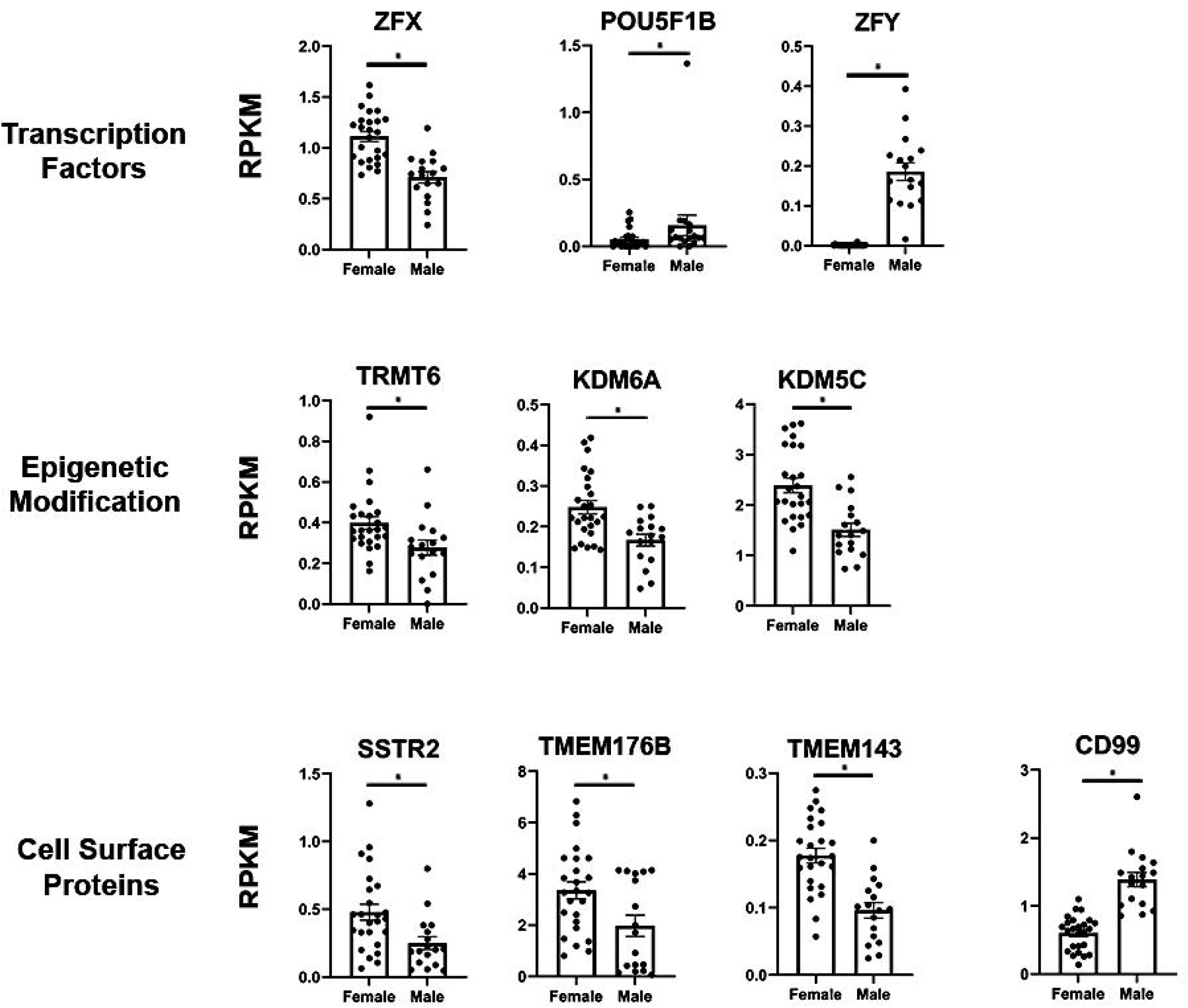
Sexually dimorphic genes in human astrocytes. Selected genes that are significantly associated with sex, including genes encoding transcription factors, epigenetic modifying enzymes, and proteins localized to the plasma membrane.

## Discussion

We generated transcriptomic data of over 40 samples of acutely purified human astrocytes. These samples vary in age, sex, and disease state, allowing us to analyze these features in humans for the first time with RNAseq. Overall, genes associated with synaptic function change in multiple conditions, highlighting dynamism in astrocyte-synapse interactions in humans. We found that human astrocytes undergo substantial change in the tumor microenvironment, notably a reduction in genes critical for maintaining excitation-inhibition balance at synapses, while astrocytes in epileptic regions in FCD patients hardly show signs of change compared to astrocytes in control regions. We also identified changes in astrocytes across maturation, from immature fetal stages to fully mature stages. Comparing human gene expression data with data from mice demonstrated broad overlap in pathways associated with astrocyte maturation, even though there were substantial differences in the specific gene signatures. Age-associate gene expression included reduced expression of synaptic genes and upregulation of reactive markers. Finally, we discovered some of the first sexually dimorphic gene expression in human astrocytes, including several transcription factors and epigenetic factors. Together, this data elucidates several fundamental aspects of human astrocyte biology in health and disease, as well as drawing important comparisons to murine astrocytes.

### Molecular Profile of Astrocytes in Human Disease

Prior to the advent of the immunopanning technique, astrocyte purification mainly relied on serum-selection (McCarthy and de Vellis, 1980). In these methods, heterogenous collections of cells were cultured with serum-containing media that preferentially allowed survival and propagation of astrocytes. However, these conditions were not physiological, as serum is a component of the blood that does not cross the blood brain barrier in healthy brain tissue. Astrocytes placed under these conditions upregulate reactive markers and adopt fibroblast-like morphology (Foo et al., 2011; Zamanian et al., 2012). Using this method, it was challenging to study *in vivo* reactive astrocyte states, as the signal was masked by the response to serum during *in vitro* purification. Immunopanning allows for acute purification without the use of cell culture or serum, maintaining astrocytes in a near-physiological state (Zhang et al., 2016). This technique allowed us to characterize the transcriptomic profile of human astrocytes from two *in vivo* neurological disorders, epilepsy and brain tumor.

Despite previous evidence that some forms of epilepsy can induce astrocyte reactivity (Binder and Steinhauser, 2006), our analysis does not find notable changes in the astrocytes collected for this study. This may reflect differences in disease progression across different kinds of epilepsy, as this dataset specifically utilizes tissue from patients with a specific form of developmental epilepsy (FCD). A human study of patients with FCD only observed astrocyte reactivity in the center of the disorganized cortex, not in outer regions with milder neuronal phenotypes (Rossini et al., 2017). Therefore, it is conceivable that human astrocytes in this epileptic context would not demonstrate reactivity.

In stark contrast, peritumor astrocytes demonstrate a robust change in gene expression. Peritumor astrocytes strongly upregulate genes related to cell cycle and proliferation, which is consistent with astrocyte reactivity. A variety of tumor cells are known to promote astrocyte reactivity and proliferation via secreted molecules, such as RANKL, in mouse models (Kim et al., 2014). Peritumor astrocytes also downregulate genes involved in a variety of astrocytic functions, including blood circulation, extracellular matrix composition, and synaptic function. Among the plethora of extracellular matrix genes that change expression, peritumor astrocytes upregulate production of matrix metallopeptidase 2 (MMP2), which degrades components of the extracellular matrix and can aid in cell invasion (Dong et al., 2011). Peritumor astrocytes strongly decrease expression of glutamate transporters (SLC1A2 and SLC1A3) that are normally highly expressed in astrocytes and help maintain the excitation-inhibition balance of the brain. Seizure activity is common in individuals with brain tumors, and seizures are often the precipitating event that leads to medical treatment (Englot et al., 2016). Astrocytes may contribute to unregulated excitation in the brain by downregulating these important glutamate transporters. Excessive excitation not only causes harmful seizures; there is recent evidence that neurons form synaptic structures with tumor cells, and neuronal activity drives further proliferation and infiltration of tumor cells (Venkataramani et al., 2019; Venkatesh et al., 2019; Zeng et al., 2019). This raises the exciting possibility of therapeutically targeting glutamate uptake in astrocytes, in addition to existing anti-seizure medications. In future studies, it will be vital to determine whether these general patterns hold for the various kinds of brain tumors, and at what stage of disease progression they appear. Beyond glutamate transporters, peritumor astrocytes downregulate several other genes that normally support synaptic function, suggesting further impacts on circuit function. Of note, there is a previously published analysis of human astrocytes inside the tumor core, as opposed to the peritumor regions examined in this study. These researchers found upregulation of proliferative markers as well, but intriguingly, they detected changes in the JAK/STAT and interferon gamma response, which is not present in the peritumor astrocytes (Henrik Heiland et al., 2019). These findings may reflect differing properties and functions of astrocytes in the tumor microenvironment and the tumor core.

### Human Astrocyte Maturation

Human brain development proceeds through a cascade of complex and reciprocal interactions between several maturing cell types. For example, neurons largely fail to make functional synapses in the absence of astrocytes, and astrocytes lack morphological complexity without the presence of neurons (Banker, 1980; Stogsdill et al., 2017). The developmental trajectory of most major brain cells has been described by the presence of cell-specific transcription factors that drive cells toward a specific fate, such as NEUROD1 in neurons and OLIG2 in oligodendrocytes. Although some important regulators have been identified, astrocyte development and maturation remain less well understood (Kang et al., 2012; Glasgow et al., 2014; Chaboub et al., 2016; Li et al., 2019). Previous work compared fetal astrocytes to postnatal astrocytes that helped identify markers of mature versus immature astrocytes, and here we extend that work by creating a developmental timeline across postnatal astrocyte maturation in humans. This provides some of the first insight into how human astrocytes mature past the early stages of development. Synaptic genes do not show differences in expression in the postnatal epoch, though they show strong upward trends before birth. What we found instead was a shift in astrocytic gene expression around 8 years old that persists into early adulthood. This time frame coincides with the onset of increased synaptic pruning in the cortex, as evidenced by a decline in cortical synaptic spine density beginning around puberty (Huttenlocher, 1979; Huttenlocher and Dabholkar, 1997; Petanjek et al., 2011). Based on this data, astrocytes adopt functions for the support and maintenance of synapses before birth, but they adopt new roles in synaptic remodeling during postnatal maturation. These data also provide new markers to use in untangling astrocytic gene networks and molecular mechanisms that related to increased synaptic pruning. Pathway analysis of astrocyte maturation genes identified downregulation of proliferative pathways and upregulation of pathways related to ion homeostasis and lipid metabolism. We also find these patterns of astrocyte maturation preserved in mice, even though mice and humans showed divergent sets of maturation-related genes. This suggests mice and human astrocytes share many aspects of their developmental arcs, though they may express different sets of genes to achieve the same functional goal. Future studies should aim to identify the signaling mechanisms that drive astrocyte maturation, as aberrations in astrocyte maturation could contribute to dysfunction in neural circuits and ultimately neurodevelopmental disorders.

### Aging in Human Astrocytes

Astrocytes become reactive in age-related neurological diseases such as Alzheimer Disease (Beach et al., 1989). It is important to determine whether reactivity is induced purely by disease progression or whether astrocyte reactivity occurs in the course of normal aging, which may further contribute to aspects of disease progression. We observe declining expression of synaptic genes, including the glutamate transporter gene SLC1A3. As we noted in peritumor astrocytes, altered expression of this gene product would impact the balance of excitation-inhibition in the brain and impair circuit function. Two separate animal studies have identified increasing expression of reactive markers across age in the mouse brain, both in the cortex and subcortical regions (Boisvert et al., 2018; Clarke et al., 2018). We find corresponding evidence in our analysis of aging human astrocytes where several genes involved in cytokine signaling are upregulated, including CCL2, IL6, and SOCS3. We also observe a modest increase in senescence markers CDKN1A and CDKN2A. As our cohort only extends to age 65, further study of astrocytes at more advanced ages could uncover much larger changes in the astrocyte transcriptome throughout the aging process. While reactive and senescence markers increase, we observe a decrease in genes related to energy metabolism. Astrocytes typically provide metabolic support to aid in proper neuronal function and signaling, so these changes may contribute to age-related declines in cognition. An important question for further examination is whether a decrease in neuronal support is reflective of astrocytic dysfunction or declining demand from neurons.

### Sexual Dimorphism of Human Astrocytes

Many neurological diseases have differences in incidence and prognosis depending on sex, but little is known about the mechanisms that underlie these differences. Recently, multiple findings identified sex differences in microglia, but sex differences in astrocytes remain elusive despite their extensive interactions with microglia. Slight differences in astrocyte number and morphology were reported in sub-cortical regions of the brain in rats, such as the amygdala (Mong and McCarthy, 2002; Johnson et al., 2008). Here, we report the first evidence of sexual dimorphism in human astrocytes, to the best of our knowledge. Female cortical astrocytes have higher transcription of several plasma membrane proteins, including somatostatin receptor SSTR2, a transcriptional target of p53, PERP, and transmembrane protein TMEM176B. We also observe differential expression of genes with epigenetic functions, such as demethylases KDM5C and KDM6A. Though healthy astrocytes demonstrate relatively few sex differences, further studies should investigate whether underlying differences in epigenetic state could contribute to sex-specific responses to insult or injury and ultimately underlie sex differences in neurological disease.

As we continue to identify astrocytic roles in human health and disease, it is imperative that we expand our knowledge of astrocyte biology, especially in human contexts. These human transcriptomic profiles are an important step in that direction, and they will provide valuable insight for further investigation of human biology and novel approaches for neurological disease.

## Acknowledgements

We thank Michael Sofroniew, Baljit Khakh, and Jessica Rexach for advice. We thank the Eli and Edythe Broad Center of Regenerative Medicine and Stem Cell Research, UCLA BioSequencing Core Facility for their services. This work is supported by the Achievement Rewards for College Scientists Foundation Los Angeles Founder Chapter and the National Institute of Mental Health of the National Institutes of Health (NIH) Award T32MH073526 to M.C.K., the Neurosurgery Research and Education Foundation Kate Carney Family Young Clinician Investigator Award to A.C.W., the National Institute of Neurological Disorders and Stroke of the National Institute of Health (NIH) R00NS089780, R01NS109025, the National Institute of Aging of the NIH R03AG065772, National Center for Advancing Translational Science UCLA CTSI Grant UL1TR001881, the W. M. Keck Foundation Junior Faculty Award, UCLA Eli and Edythe Broad Center of Regenerative Medicine and Stem Cell Research Innovation Award, and the Friends of the Semel Institute for Neuroscience & Human Behavior Friends Scholar Award to Y. Z.

